# Functional variability in adhesion and flocculation of yeast megasatellite genes

**DOI:** 10.1101/2022.01.14.476295

**Authors:** Cyril Saguez, David Viterbo, Stéphane Descorps-Declère, Brendan Cormack, Bernard Dujon, Guy-Franck Richard

## Abstract

Megasatellites are large tandem repeats found in all fungal genomes but especially abundant in the opportunistic pathogen *Candida glabrata*. They are encoded in genes involved in cell-cell interactions, either between yeasts or between yeast and human cells. In the present work, we have been using an iterative genetic system to delete several *C. glabrata* megasatellite-containing genes and found that two of them were positively involved in adhesion to epithelial cells, whereas three genes controlled negatively adhesion. Two of the latter, *CAGL0B05061g* or *CAGL0A04851g,* are also negative regulators of yeast-to-yeast adhesion, making them central players in controlling *C. glabrata* adherence properties. Using a series of synthetic *Saccharomyces cerevisiae* strains in which the *FLO1* megasatellite was replaced by other tandem repeats of similar length but different sequences, we showed that the capacity of a strain to flocculate in liquid culture was unrelated to its capacity to adhere to epithelial cells or to invade agar. Finally, in order to understand how megasatellites were initially created and subsequently expanded, an experimental evolution system was set up, in which modified yeast strains containing different megasatellite seeds were grown in bioreactors for more than 200 generations and selected for their ability to sediment at the bottom of the culture tube. Several flocculation-positive mutants were isolated. Functionally relevant mutations included general transcription factors as well as a 230 kb segmental duplication.

## INTRODUCTION

All eukaryotic genomes sequenced so far contain a variable amount of tandemly repeat DNA sequences (Richard *et al*. 2008). These can be classified in three main categories, according to the length of their structural motif. Microsatellites are made of 1-9 bp motifs and are very abundant in all genomes. The *S. cerevisiae* genome contains 1818 di-, tri- and tetranucleotide repeats, the three most abundant microsatellites (Malpertuy *et al*. 2003), whereas the human genome contains 260,000 such repeats per haplotype (International Human Genome Sequencing Consortium 2001). Minisatellites are made of slightly larger motifs (10-90 bp) and are less frequent. Less than 100 such tandem repeats were found in the budding yeast genome, mainly in genes encoding cell wall proteins (Bowen *et al*. 2005; Verstrepen *et al*. 2005; Richard and Dujon 2006). Finally, an additional family was proposed to encompass tandem repeats whose structural motif was larger, those were called megasatellites (Thierry *et al*. 2008, 2009). Initially described in the pathogenic yeast *Candida glabrata* as tandem repeats whose base motif was larger than 100 bp, this was refined by a subsequent study covering 21 fungal genomes, in which it was found that length distribution of minisatellites and megasatellites were different. It was therefore chosen to use a length cutoff between the two kinds of tandem repeats of 90 bp, with those whose structural motif was larger being called megasatellites (Figure 1A) (Tekaia *et al*. 2013).

**Figure 1:**
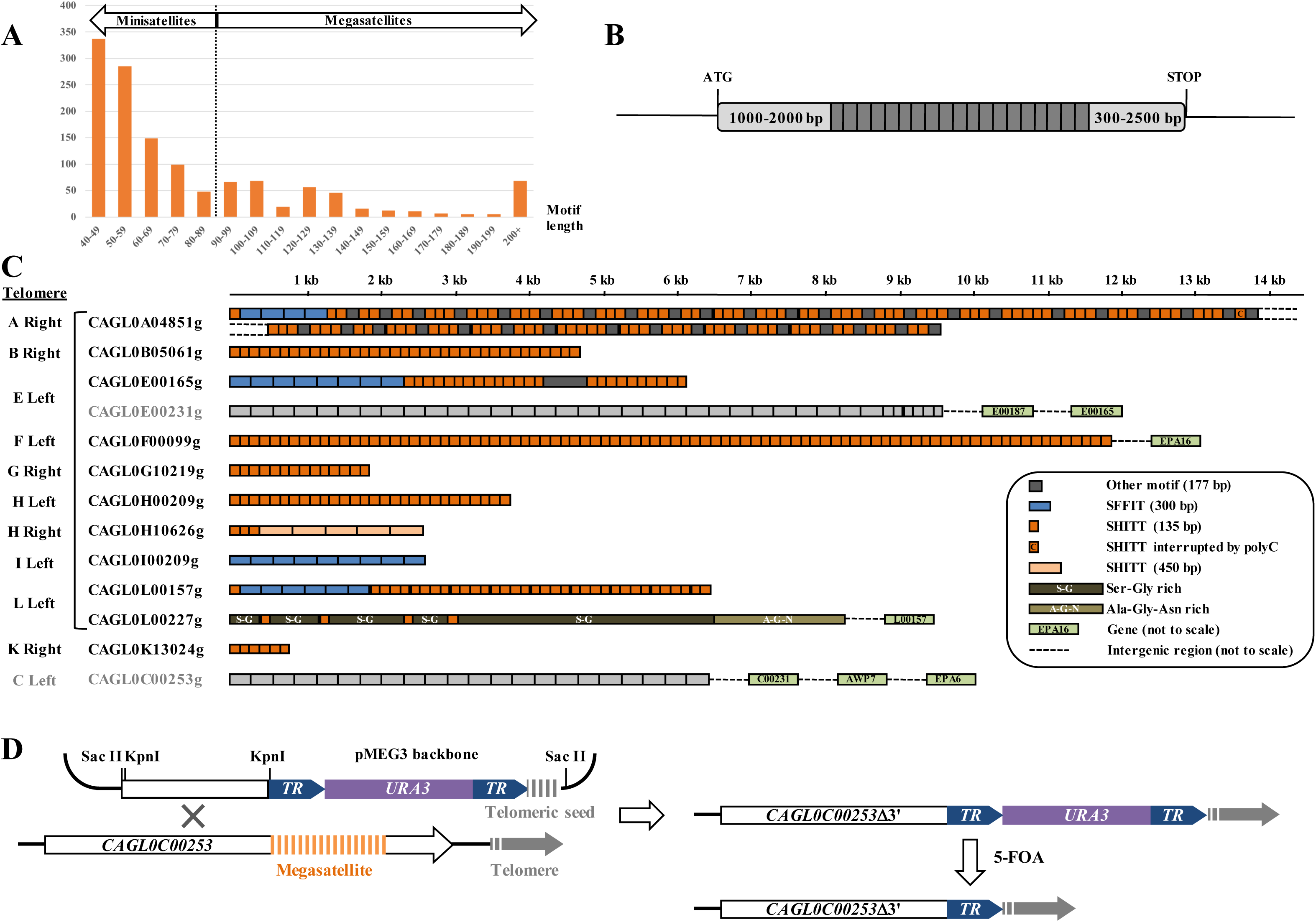
Megasatellite distribution in *Candida glabrata*. **A**: Tandem repeats detected in 21 fungal genomes were plotted as a function of motif length (Tekaia *et al*. 2013). The transition from minisatellites to megasatellites was set around 90 bp (dotted line). Note that minisatellites whose motif was shorter than 40 bp were not shown for the sake of clarity. **B**: General organization of a megasatellite-containing gene. The 5’ end of the gene, before the tandem repeat, is usually 1-2 kb, while the 3’ end exhibits more length variability. Motifs are always in frame and their number ranges from 3 to 30 in *C. glabrata*. **C**: Telomeric megasatellites in *C. glabrata*. Chromosome numbers are indicated to the left, along with the telomeric arm. The bracket includes all the genes part of a large paralogous family. SHITT and SFFIT motifs are indicated by a color code, orange or blue respectively. Motifs in dark grey are intervening sequences. *CAGL0L00227g* contains large regions almost entirely made of glycine and serine residues (S-G) or alanine, glycine and asparagine residues (A-G-N), extending sometimes over considerable distances. Note that two megasatellite-containing genes (*CAGL0E00231g* and *CAGL0C00253g*) could not be deleted despite repetitive attempts and are therefore indicated in light grey. All megasatellite-containing genes shown here are the last gene before the telomere, except in four cases (*CAGL0E00231g*, *CAGL0F00099g*, *CAGL0L00227g* and *CAGL0C00253g*). In these four cases, the gene(s) between the megasatellite and the telomere are indicated. All drawings are oriented in such a way that telomeric ends are on the right. **D**: Experimental setup to delete telomeric megasatellites. A PCR product containing 1 kb of DNA upstream the repeat tract was cloned at the *Kpn*I site of pMEG3. The resulting plasmid was then linearized with *Sac*II and transformed into *C. glabrata*. Homologous recombination with the telomeric sequence led to the deletion of all sequences downstream the PCR product and addition of a new telomere on the telomeric seed. Spontaneous single-strand annealing events between the two tandem repeats (*TR*, in blue) were selected on 5-FOA medium, so that the resulting strain could be deleted again with *URA3*. Up to 11 megasatellites were iteratively deleted using this approach.

*Candida glabrata* contains 44 megasatellites, in 33 different genes (Thierry *et al*. 2008). Most of the protein functions encoded by these genes are unknown, but many carry signatures of cell wall proteins and are good candidates to be involved in yeast adhesion to epithelial cells. The structure of megasatellite-containing genes is always the same: the tandem repeat is located in the middle of the gene, 1-2 kb after the start codon and 300-2500 bp from the stop codon, always in frame (Figure 1B). Several families of motifs were found to be encoded by megasatellites, but two were particularly frequent and were called SHITT and SFFIT motifs, based on the eponymous five amino acids conserved in the translation products of all motifs of the family (Figure 1C). The duplication and evolution of these motifs was studied by clustering analyses and showed recurrent transfer of genetic information between megasatellites (Rolland *et al*. 2010). Recent resequencing of *C. glabrata,* using a mix of long and short reads, allowed to make substantial corrections to the reference genome. Forty-five genes that were misassembled were removed or fixed, 31 new open reading frames were annotated and 21 repeat-containing genes were corrected, establishing a new high quality reference for the *C. glabrata* genome (Xu *et al*.). By chromosomal conformation capture, there was no evidence that megasatellites cluster within the *C. glabrata* nucleus, nor that they were frequently associated with replication origins or terminations (Descorps-Declère *et al*. 2015). *S. cerevisiae* also contains several megasatellites, although fewer than its pathogenic cousin. The best known gene family containing megasatellites is the FLO family, in which the *FLO1*, *FLO5* and *FLO9* genes encode a 135 bp FLO motif, rich in threonine residues (Richard and Dujon 2006; Rolland *et al*. 2010). *FLO1* has been identified for a long time as being one of the genes responsible for budding yeast flocculation in liquid culture and the length of the FLO megasatellite was shown to be positively correlated to the extent of flocculation (Verstrepen *et al*. 2005). Other yeast species also contain megasatellites, the most widespread motif being related to the FLO motif (Tekaia *et al*. 2013). *Candida albicans* and *Candida dubliniensis* genomes contain several ALS megasatellites encoded by the eponymous adhesin gene family, involved in yeast adhesion to epithelial cells (Hoyer 2001). Interestingly, very recent work showed that megasatellite length in two *S. cerevisiae* genes (*HPF1* and *FLO11*) is correlated with life span, as determined by QTL analyses. Repeat expansion in the *HPF1* gene shifted yeast cells from a sedimenting to a buoyant state, completely modifying oxygenation as well as the surrounding metabolism, resulting in shorter life span (Barre *et al*. 2019).

The aim of the present work was to decipher the function of megasatellite-containing genes in *Candida glabrata*, as well as setting up an experimental evolution assay, using *FLO1*-dependent flocculation, to catch primary events leading to megasatellite formation in *Saccharomyces cerevisiae*. By iteratively deleting *C. glabrata* subtelomeric megasatellites, we found that some deletions increased cellular adhesion whereas others decreased it, suggesting a complex role for megasatellite-containing genes. The experimental evolution assay allowed the isolation of flocculation mutants, but none of them showed an amplification of the *FLO1* megasatellite. Using synthetic *FLO1* genes, we found that different tandem repeats play distinct roles in flocculation and cell-to-cell adhesion.

## MATERIALS AND METHODS

### *C. glabrata* plasmids

A 578 bp piece of *Yarrowia lipolytica* genome located in an intergenic region (Chrom. A, 2057200-2057778) was PCR amplified using primers YALIupfor and YALIuprev (Supplemental Table 1, top). The PCR product was digested with *Kpn*I and *Bam*HI and cloned in pBlueScript SK+ at the corresponding sites. The resulting plasmid (pMEG0) was digested with *Bgl*II and *Bam*HI and ligated to the *C. glabrata URA3* gene amplified using CgURA3 primers and digested with the same restriction enzymes, to give plasmid pMEG1. The same 578 bp piece of *Y. lipolytica* genome was amplified with YALIdownfor and YALIdownrev primers, digested with *Bgl*II and *Not*I and cloned into pMEG1 at the corresponding restriction sites. The resulting plasmid (pMEG2) was subsequently digested with *Not*I, dephosphorylated and ligated to phosphorylated TELup and TELdown complementary oligonucleotides, encoding four *C. glabrata* telomeric repeats (Kachouri-Lafond *et al*. 2009), to give plasmid pMEG3. All subsequent constructs used to delete telomeric megasatellites were made in pMEG3. For each deletion, ca. 1000 bp upstream to the megasatellite were amplified using dedicated primers (Supplemental Table 1, bottom). PCR products were digested by *Kpn*I and cloned into pMEG3 at the *Kpn*I site, to give plasmids pMEG4 to pMEG18. Note that some PCR products could not be cloned, therefore initially planned deletions of genes *CAGL0E00231g* and *CAGL0C00253g*, were not achieved (Figure 1C, genes in grey).

### *S. cerevisiae* plasmids

Synthetic FLO genes were assembled in the pRS406 integrative plasmid, carrying *URA3* as a selection marker (Sikorski and Hieter 1989). pRS406-*FLO1*ΔR was built as follows: primers SC1 and SC6 were used to amplify a 1272 bp DNA fragment from the *FLO1* gene upstream the megasatellite and overlapping pRS406; primers SC4 and SC5 were used to amplify a 1776 bp DNA fragment from the *FLO1* gene downstream the megasatellite and overlapping pRS406 (Supplemental Table 2). Both PCR products were assembled into pRS406 using Gibson assembly mix (NEBiolabs). Plasmid pRS406-*FLO1*ΔR::FLO was built as follows: primers SC1 and SC2 were used to amplify a 1252 bp DNA fragment from the *FLO1* gene upstream the megasatellite and overlapping pRS406; primers SC3 and SC4 mlwere used to amplify a 1753 bp DNA fragment from the *FLO1* gene downstream the megasatellite and overlapping pRS406. Both PCR products were assembled into pRS406 along with primers SC7, SC8, SC9 and SC10 using Gibson assembly. This reconstituted a *FLO1* gene containing only one FLO motif. Plasmid pRS406-*FLO1*ΔR::SHITT was built the same way, except that primers SC11, SC12, SC13 and SC14 were used in the Gibson assembly to reconstitute a *FLO1* gene containing only one SHITT motif. Plasmid pRS406-*FLO1*ΔR::ALS was built the same way, except that primers SC15, SC16, SC17 and SC18 were used in the Gibson assembly to reconstitute a *FLO1* gene containing only one ALS motif. Plasmid pRS406-*FLO1*ΔR::2FLO was built as follows: primers SC8, SC9, SC10 and SC19 were phosphorylated and ligated *in vitro* using T4 DNA ligase to make a 178 bp piece of DNA containing one FLO motif. primers SC7, SC8, SC9 and SC20 were phosphorylated and ligated to make a 175 bp piece of DNA also containing one FLO motif. Both FLO motifs were ligated and gel purified to make a 353 bp piece of DNA containing two FLO motifs. This DNA was assembled into pRS406 along with SC1-SC2 and SC3-SC4 PCR products using Gibson assembly. This reconstituted a *FLO1* gene containing a tandem of two FLO motifs.

The *FLO1* genes containing synthetic tandem repeats were designed and assembled at ProteoGenix. They were delivered as identical copies of *FLO1* containing 10 tandemly repeated FLO motifs, 10 FLOamy motifs, 10 SHITT motifs or 13 ALS motifs, inserted at the normal location within the gene. Synthetic *FLO1* genes were delivered cloned in pUC57 and were transferred in pRS406 for further integration in yeast cells.

### *Candida glabrata* strains

All megasatellite deletions were made in the HM100 strain, a derivative of the reference CBS138 strain, or in the BG14 strain, a derivative of the commonly used BG2 strain. Both HM100 and BG14 strains were inactivated for the *URA3* gene. Each pMEG plasmid was digested with *Sac*II in order to release the recombinogenic DNA fragment (Figure 1D) and transformed in *C. glabrata* following the lithium acetate protocol used for *S. cerevisiae* (Gietz *et al*. 1995). Transformants were subcloned on synthetic SC-Ura dropout medium before DNA extraction and molecular analysis. For each deletion, eight transformants were analyzed by Southern blot according to published methods (Viterbo *et al*. 2018). Transformants showing the expected pattern of bands were patched on yeast complete medium (YPD) and grown for 2-3 days at 30°C, before being replica plated on a 5-FOA plate supplemented with 15-20 mM nicotinamide, in order to unsilence subtelomeric regions. Without nicotinamide the whole patch was growing due to silencing of the *URA3* gene and we were unable to select [Ura-] clones. One or two [Ura-] colonies were subcloned on 5-FOA plate supplemented with nicotinamide, before PFGE analysis. All strains are described in Supplemental Table 3.

### *S. cerevisiae* strains

Total gene deletions of *FLO9* and *FLO11* were carried out by classical “ends out” recombination (Baudin *et al*. 1993). *FLO8* correction, *FLO5* and *FLO10* megasatellite deletions, as well as all modifications of the *FLO1* gene were performed by the classical two-step replacement method (Sherer and Davis 1979). Note that only the repeated part of *FLO5* and *FLO8* were deleted, the remaining portions of the gene, 5’ and 3’ of the megasatellite, were kept in frame. This is the meaning of “ΔR” alleles indicated in Supplemental Table 3. Transformants were analyzed by Southern blot to verify all constructs.

### Pulse-field gel electrophoresis of *C. glabrata* mutants

All [Ura-] mutants were analyzed by PFGE in order to check that the expected chromosome had been targeted. Yeast cells were grown to stationary phase in YPD, overnight at 30°C. In the morning, ca. 5 x 10^8^ cells were collected, centrifuged and washed with 5 mL 50 mM EDTA (pH 9.0). The pellet was resuspended in 330 μL 50 mM EDTA (pH 9.0), taking into account the pellet volume. Under a chemical hood, 110 μL of Solution I (1 M sorbitol, 10 mM EDTA (pH 9.0), 100 mM sodium citrate (pH 5.8), 2.5% β-mercaptoethanol and 10 μL of 100 mg/mL Zymolyase 100T-Seikagaku) were added to the cells, before 560 mL of 1% InCert agarose (Lonza) were delicately added and mixed. This mix was rapidly poured into plug molds and left in the cold room for at least 10 minutes. When solidified, agarose plugs were removed from the molds and incubated overnight at 37°C in Solution II (450 mM EDTA (pH 9.0), 10 mM Tris-HCl (pH 8.0), 7.5% β-mercaptoethanol). In the morning, tubes were cooled down on ice before Solution II was delicately removed with a pipette and replaced by Solution III (450 mM EDTA (pH 9.0), 10 mM Tris-HCl (pH 8.0), 1% N-lauryl sarcosyl, 1 mg/mL Proteinase K). Tubes were incubated overnight at 65°C, before being cooled down on ice in the morning. Solution III was removed and replaced by 500 mM EDTA (pH 9.0) before being loaded on gel. A 1% SeaKem agarose gel (Lonza) was poured in 0.25 X TBE buffer, plugs were loaded and the run was performed on a Rotaphor machine (Biometra) in 0.25 X TBE. Parameters chosen for *C. glabrata* chromosomes were set on: initial pulse: 200 seconds, final pulse: 70 seconds, run time: 70 hours, voltage: 140 V, angle: 120° (linear), temperature: 12°C. At the end of the run, the gel was stained in ethidium bromide, before being transfered for hybridization, as previously described (Viterbo *et al*. 2018).

### Evolution to flocculation experiment settings

Cells were grown in 55 mL bioreactors, in rich medium (YPD) at 30°C under constant oxygenation. The oxygen inlet reached the bottom of the tube, therefore flocculating cells would obtain a slight growth advantage over buoyant cells. Preliminary tests showed that flocculating yeasts were more frequent when cells were grown to stationary phase rather than when continuously grown in exponential phase. Stationary phase was reached around OD 25 (1.8 x 10^8^ cells/mL). After one week, 50 mL of culture was removed from the top of the bioreactor. The bottom 5 mL was homogenized and diluted to OD ∼0.1 (7 x 10^5^ cells/mL), in fresh YPD, after which cells started to grow exponentially until reaching stationary phase (Supplemental Figure 1). This was performed 27 times in a row over a period of six months, resulting in a total of 214-218 generations for each strain. When the whole culture was entirely flocculating, it was isolated and analyzed. A fresh culture was restarted from the frozen stock (generation 0). The [Flo+] phenotype was confirmed by a larger culture in flask allowing to assess the presence of large flocs each containing millions of yeast cells. Calcium-dependence flocculation phenotype was first verified by adding EDTA to the liquid culture, before precise identification of the mutation by molecular means. Each mutant was subsequently tested for dominance/recessivity and complementation tests with known flocculation mutants (or whole genome sequencing if complementation proved to be negative), as well as invasion tests on agar plates.

### Adhesion on epithelial cells

Lec2 cells were grown to confluence in 24-well microplates, fixed with 2% paraformaldehyde for 2 hours, washed four times with PBS and stored at 4°C in PBS + Pen/Strep. Yeast cells were grown in YPD to stationary phase, diluted 1/20° in YPD supplemented with 20 mM nicotinamide in order to unsilence subtelomeric regions (De Las Penas *et al*. 2003), and grown for another 3 hours at 30°C. Cultures were washed three times in 1X HBSS supplemented with 5 mM CaCl2. Cell concentration was determined and adjusted to 10^7^ cells/mL. Three dilutions of this inoculum were plated on YPD to serve as the input. Three wells of the same Lec2 cells fixed in 24-well microplate were incubated with 1 mL of the inoculum and incubated 5 minutes at room temperature. The microplate was spined down at 100 rpm for one minute, then incubated at room temperature for 10 minutes. The microplate was inverted to remove inocula and each well was washed four times with 500 μL HBSS supplemented with 5 mM CaCl2. Finally, 500 μL of cell lysis buffer (1X PBS, 10 mM EDTA (pH 8.0), 0.05% Triton X100) was added to each well, cells were scraped thoroughly, diluted to an appropriate concentration and plated on YPD to serve as the output. Adhesion was calculated as the ratio of output CFU/input CFU. Note that given the experimental variability from plate to plate, an appropriate wild-type control (CBS138 or BG2) was added in triplicate in each plate.

### Invasion of agar plates

Yeast cells were grown to stationary phase in YPD, then patched on YPD and grown for 6-10 days at 30°C. Plates were gently washed under running water until the cell layer was removed and plates were incubated an extra 24 hours at 30°C. Adhesion was visually evaluated according to the amount of growth visible after 24 hours and classified in three categories: no adhesion, weak adhesion, strong adhesion.

### Flocculation tests

Yeast cells were grown to stationary phase in YPD. Culture tubes were vortexed and left one minute to stand on the bench before 200 μL were collected right below the meniscus. Optical density at 600 nm of collected cells was determined and used as a proxy for flocculation capacity.

### Spheroplast rate assay

For each of the two yeast species, a lysis curve was first established with wild-type strains (BY4741 and HM100), as previously published (Ovalle *et al*. 1998). Cells were grown overnight to stationary phase in YPD and diluted 1/50° in the morning in 50 mL fresh medium. When cell concentration reached 10^7^-3 x 10^7^ cells/mL, 3 x 10^8^ cells were collected and washed thrice with sterile water in a 50mL polypropylene tube. Cells were resuspended in 15 mL 10 mM Tris buffer pH 8.0, 10 mM EDTA pH 8.0, to disrupt potential flocculation aggregates. Zymolyase (100T, Seikagaku) was added at a final concentration of 3.3 μg/mL. The tube was incubated at 25°C and 1 mL of cell suspension (2 x 10^7^ cells ) was collected every 5 minutes during 75 minutes. Cells were diluted 1/10° in water and optical density at 600 nm was determined. OD were plotted at each time point to identify the linear part of the lysis curve (Ovalle *et al*. 1998). This allowed to determine that in subsequent experiments with *S. cerevisiae* and *C. glabrata* mutants, OD measurements should be performed after 10 minutes of incubation with zymolyase.

For each mutant and wild type controls, yeast cells were grown overnight to stationary phase, in 3 mL YPD at 30°C. In the morning, 1/50° dilutions were performed in 3 mL fresh YPD. When cell concentration reached 10^7^-3 x 10^7^ cells/mL, 2 x 10^7^ cells were collected, washed thrice with sterile water in a microtube, and resuspended in 1 mL 10 mM Tris buffer pH 8.0, 10 mM EDTA pH 8.0. Zymolyase was added at a final concentration of 3.3 μg/mL and tubes were incubated at 25°C for 10 minutes, before being transferred on ice to stop the reaction. After pipetting several times to homogenize, 100 μL cells were collected, 900 μL water were added and OD at 600 nM was determined for each sample tube. All experiments were performed 3 times for each strain.

All statistical analyses were performed using the ’R’ package (Millot 2011).

All reagents, plasmids and yeast strains described in the present manuscript are freely available on request.

## RESULTS

### Megasatellite-containing genes play different roles in *C. glabrata* cellular adhesion

In a first series of experiments, we intended to delete all telomeric megasatellites, using a reusable marker strategy and a telomeric seed sequence (Sandell and Zakian 1993). The upstream sequence of each gene was PCR amplified and cloned in a plasmid (pMEG3, Figure 1D) containing the *C. glabrata URA3* gene flanked by a 600 bp tandem repeat sequence from *Yarrowia lipolytica* (*TR*, Figure 1D). After linearization, each plasmid was transformed into *C. glabrata* and [Ura+] transformants were recovered. All of them were analyzed by Southern blot, in order to verify that the integration site was correct. Each *bona fide* deletant was then plated on 5-FOA medium in order to select [Ura-] subclones in which the *URA3* marker had been lost by recombination between the two *TR* sequences. Following this procedure, resulting strains could then be reused to delete a second telomeric megasatellite. Up to eleven megasatellites were iteratively deleted in the same strain using this approach.

All clones were analyzed by Southern blot, using either the *URA3* gene or the TR repeat as a probe. Hybridization with the *URA3* probe revealed a band whose molecular weight was 2830 bp plus the length of the added telomeric repeat. We found that such signals could reach more than 10 kb, depending on the transformant (Figure 2A, top). However, after 5-FOA selection, telomere length decreased to a more reasonable 200-600 bp (Figure 2A, bottom). It is possible that tandem integration of the *TR*-*URA3*-*TR* cassette produced long telomeric fragments, that were lost by recombination after 5-FOA selection. Strain karyotype was also analyzed by pulse-field gel electrophoresis (PFGE), in order to separate all yeast chromosomes. The gel was thereafter transferred and hybridized using the *TR* repeat as a probe. Chromosomes carrying at least one such repeat were revealed (Figure 2B, top). In most cases, the PFGE profile perfectly matched the expected profile according to the published sequence (Figure 2B, bottom). This shows that the megasatellite genes we attempted to delete were properly assembled on their cognate chromosomes in the genome sequence (Dujon *et al*. 2004). In one instance, the targeting construct for gene *CAGL0L00157g* integrated on chromosome G instead of L. This strain was not used in subsequent experiments (CGF251 in Supplemental Table 1). All these mutants were engineered in the CBS138 type strain. When we tried to build equivalent mutations in the BG2 strain background -commonly used for adhesion studies (Cormack *et al*. 1999)- fewer transformants were obtained and most of them were not integrated at the expected locus. This is probably due to sequence polymorphisms between the two strains as well as to chromosomal translocations such as those previously described (Muller *et al*. 2009). Only one mutant could finally be built in the BG2 background (strain CGF21).

**Figure 2:**
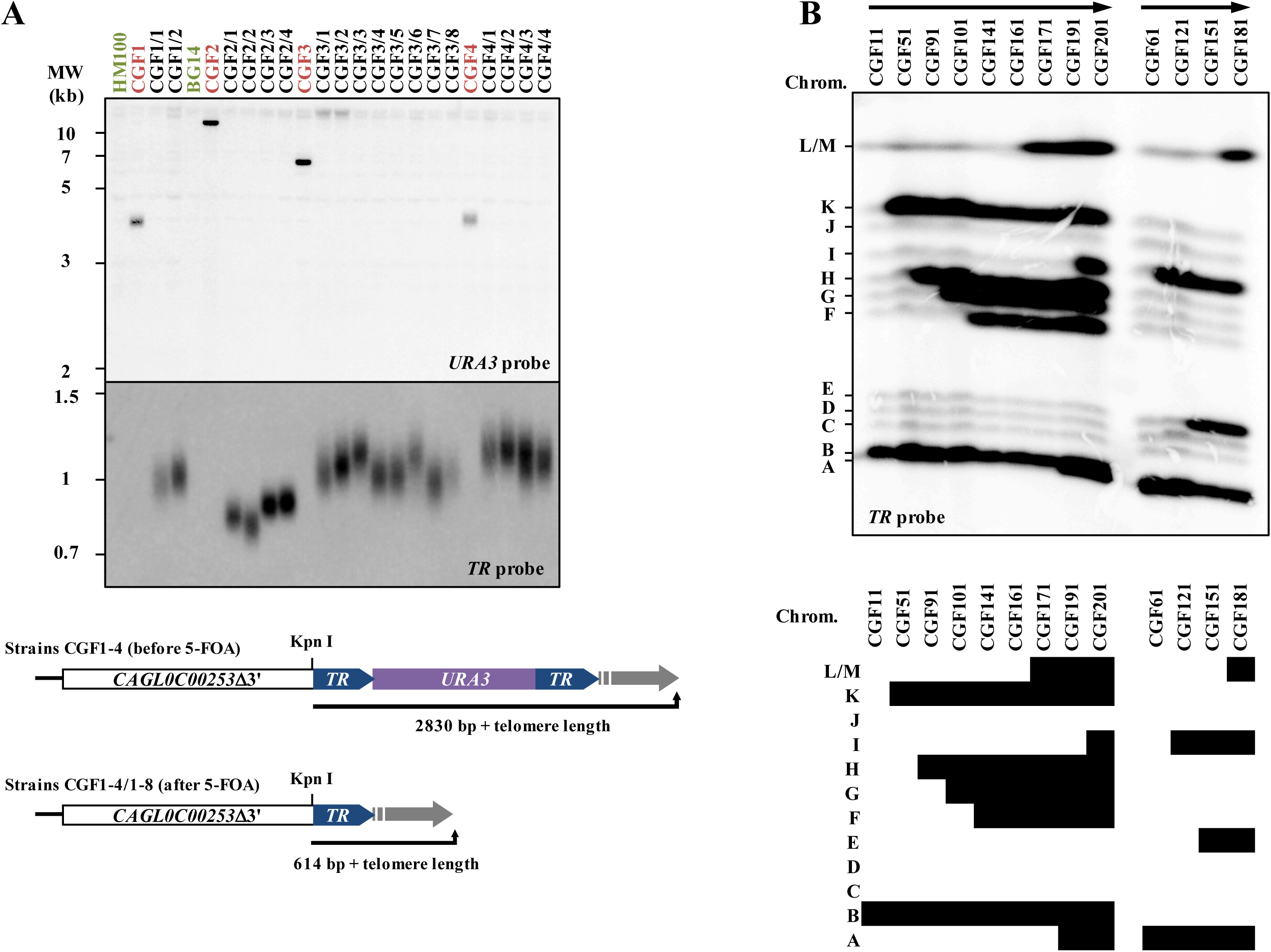
Molecular analysis of megasatellite deleted strains. **A**: Southern blot hybridized with two different probes. Wild-type controls are shown in green (CBS138 and BG2), strains after transformation but before 5-FOA selection are in red (CGF1-4), several clones of the same strains after 5-FOA selection are in black. Top: hybridization with the *URA3* probe was positive in the four *URA3*-containing strains and showed as expected different integration sites, depending on the targeted megasatellite. Bottom: hybridization with the tandem repeat (*TR*) probe shows telomere length variability in the different subclones analyzed, after 5-FOA selection. Expected molecular weights in strains before and after 5-FOA selection. **B**: Pulse-field gel electrophoresis of strains after 5-FOA selection. Top: Southern blot hybridization with the *TR* probe highlights chromosomes carrying the tandem repeat. Strains are classified from left to right in the order in which megasatellites were deleted. The cartoon below the blot depicts the expected pattern, which is strictly identical to the observed hybridization result.

We ended up generating seven single mutants and 13 multiple mutants, carrying from one to eleven subtelomeric deletions (Supplemental Table 1). All these strains were tested for their capacity to adhere to hamster epithelial cells (Lec2). (Figure 3). In each experiment, a wild-type strain (either CBS138 or BG2) served as an internal standard and the adhesion of each mutant strain was divided by the adhesion value in the standard. When this ratio was above one, the mutant was considered to adhere better than the wild type, the opposite when the ratio was lower than one. Strains deleted for *EPA1*, *EPA2*, *EPA3*, *EPA6* and *EPA7*, known adhesins in *C. glabrata*, were used as controls.

**Figure 3:**
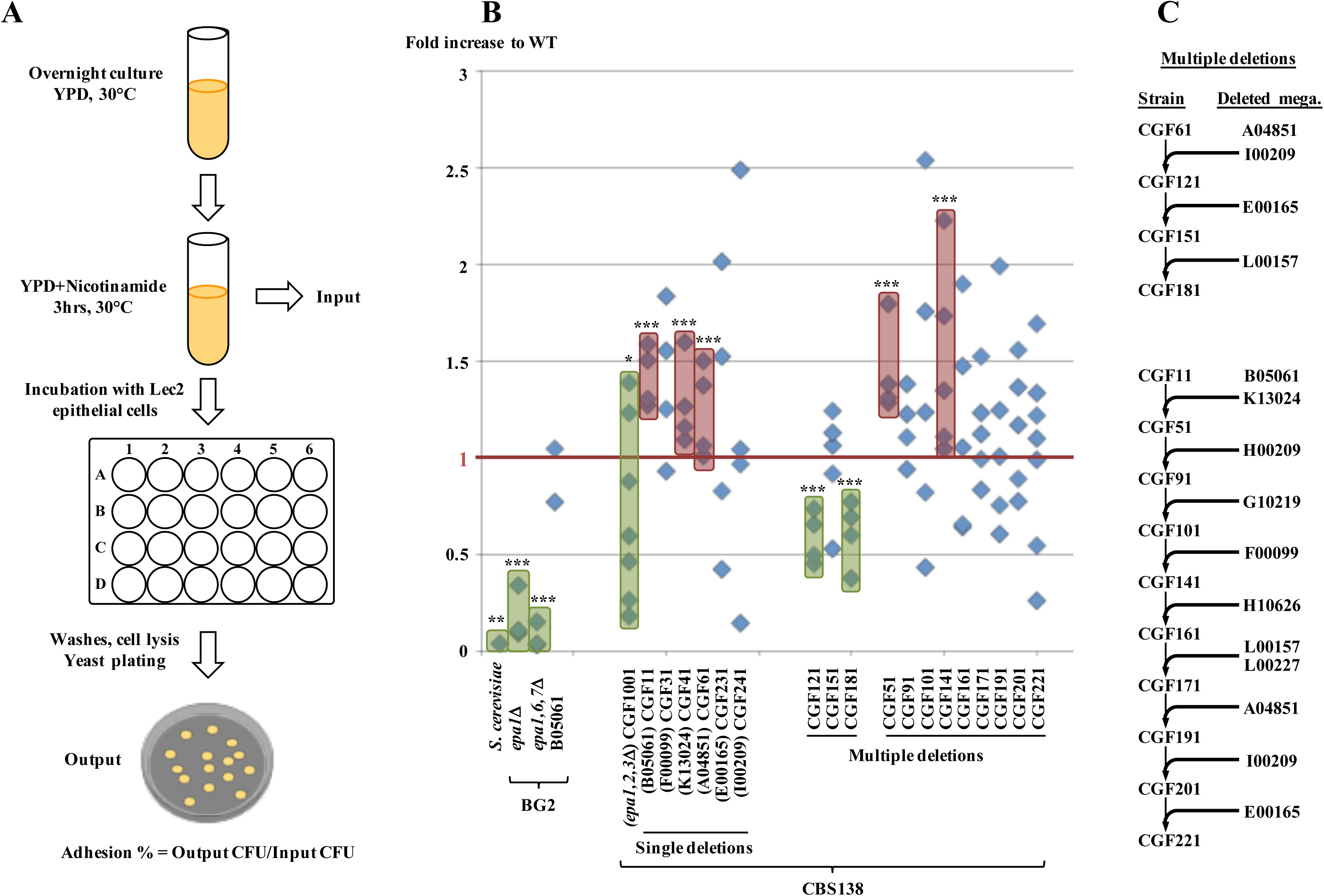
Adhesion to epithelial cells of megasatellite mutants. **A**: Experimental protocol. Adhesion values were calculated as output CFU (yeasts bound to Lec2 cells) over input CFU (yeasts before incubation with Lec2 cells). **B**: Adhesion values relative to wild type (CBS138 or BG2, depending on the mutant tested). Each diamond corresponds to one experiment, each experiment being the average of an adhesion test performed in triplicate. Note that wild type values are not included in the graph, since adhesion values in each mutant strain were divided by the adhesion value in the wild-type strain, independently determined for each experiment. Deletions that statistically increased adhesion (ratios to wild type above 1) are boxed in red, those that significantly decreased adhesion (ratios to wild type below 1) are boxed in green. Non-parametric Mann-Whitney tests were performed, asterisks showing significance levels: * p-value< 5%, ** p-value< 1%, *** p-value< 0.1%. Note that gene names are abbreviated for figure clarity, e. g. *CAGL0B05061g* was abbreviated by B05061. **C**: Summary of iterative deletions in multiple mutants. Two series of iterative deletions were made, one starting with *CAGL0A04851g* deletion, the other with *CAGL0B05061g* deletion. Note that deletion of *CAGL0L00227g* also deleted the telomere-proximal downstream gene *CAGL0L00157g*.

In the BG2 background, as expected, the *epa1*Δ mutant as well as the *epa1*Δ *epa6*Δ *epa7*Δ triple mutant were less adherent than wild type (Figure 3B). No effect was found for the deletion of *CAGL0B05061g*.

In the CBS138 background, *EPA1* (and downstream *EPA2* and *EPA3*) deletion led to reduced adhesion (Mann-Whitney test p-value= 4.1%), although markedly less than compared to the BG2 background. We concluded that *EPA1* played a more central role in adhesion in the BG2 background than in CBS138 under the conditions of this assay. Three single deletions showed a small but significant increase in adhesion: *CAGL0B05061g* (which had no effect in BG2), *CAGL0K13024g* and *CAGL0A04851g*. *CAGL0B05061g* and *CAGL0K13024g* contain pure SHITT repeats whereas *CAGL0A04851g* contains one of the longest megasatellite of the yeast genome with 113 SHITT and four SFFIT repeats (Figure 1C).

We originally expected that multiple deletions showed an additive effect on adhesion, but results were more intricate. A summary of the order in which multiple deletions were performed is shown in Figure 3C. Deleting *CAGL0I00209g* in CGF61 strain decreased adhesion, as did further deletion of *CAGL0L00157g* deletion, whereas deleting *CAGL0E00165g* did not reduce it. These three genes all contain a SFFIT repeat (Figure 1C), but in as much as deleting multiple genes did not give a simply cumulative effect on adhesion, our data do not suggest that they are functionally redundant. The next set of multiple mutations showed that deleting *CAGL0K13024g* in addition to *CAGL0B05061g* did not modify the strain ability to adhere to epithelial cells. Deletion of *CAGL0H00209g* and *CAGL0G10219g* somewhat decreased adhesion whereas deleting *CAGL0F00099g* significantly increased values above wild type in this quintuple mutant. Adding successive deletions of five more megasatellites tended to increase experimental variability, but none of these multiple mutants was statistically different from wild type. We conclude from these experiments that *CAGL0B05061g*, *CAGL0K13024g*, *CAGL0A04851g* and *CAGL0F00099g* can (at least in certain strain contexts) play a negative role in adhesion (their deletion increases adhesion) whereas *CAGL0I00209g* and *CAGL0L00157g* can play a positive role (their deletion decreases adhesion), no other gene could be shown to play any significant function. More importantly, we found no evidence that a capacity in increasing or decreasing adhesion could be specifically attributed to SFFIT or SHITT repeat-containing genes.

We subsequently performed two other assays on the same mutants: a flocculation assay to determine whether cell-to-cell adhesion between yeasts was modified and a spheroplast rate assay to determine cell wall integrity (Ovalle *et al*. 1998). The individual deletion of *CAGL0B05061g*, *CAGL0F00099g*, *CAGL0A04851g* or *CAGL0E00165g* significantly increased the ability of these mutants to flocculate (Figure 4A). When multiple deletions were examined, it was found that all strains deriving from CGF11 (deleted for *CAGL0B05061g*) also showed increased flocculation. The first conclusion that could be drawn by comparing Figures 3B and 4A is that deletion of *CAGL0B05061g* or *CAGL0A04851g* increases both adhesion to epithelial cells and between yeast cells. No other obvious correlation could be observed, showing that these two phenotypes mainly involve different genes. The second conclusion is that flocculation in multiple deletants is consistent with additional mutations not altering the phenotype. CGF121, CGF151 and CGF181 are not statistically different from each other and from the HM100 wild type reference, and the series of multiple mutations deriving from CGF51 are all significantly different from wild type (Figure 4A).

**Figure 4:**
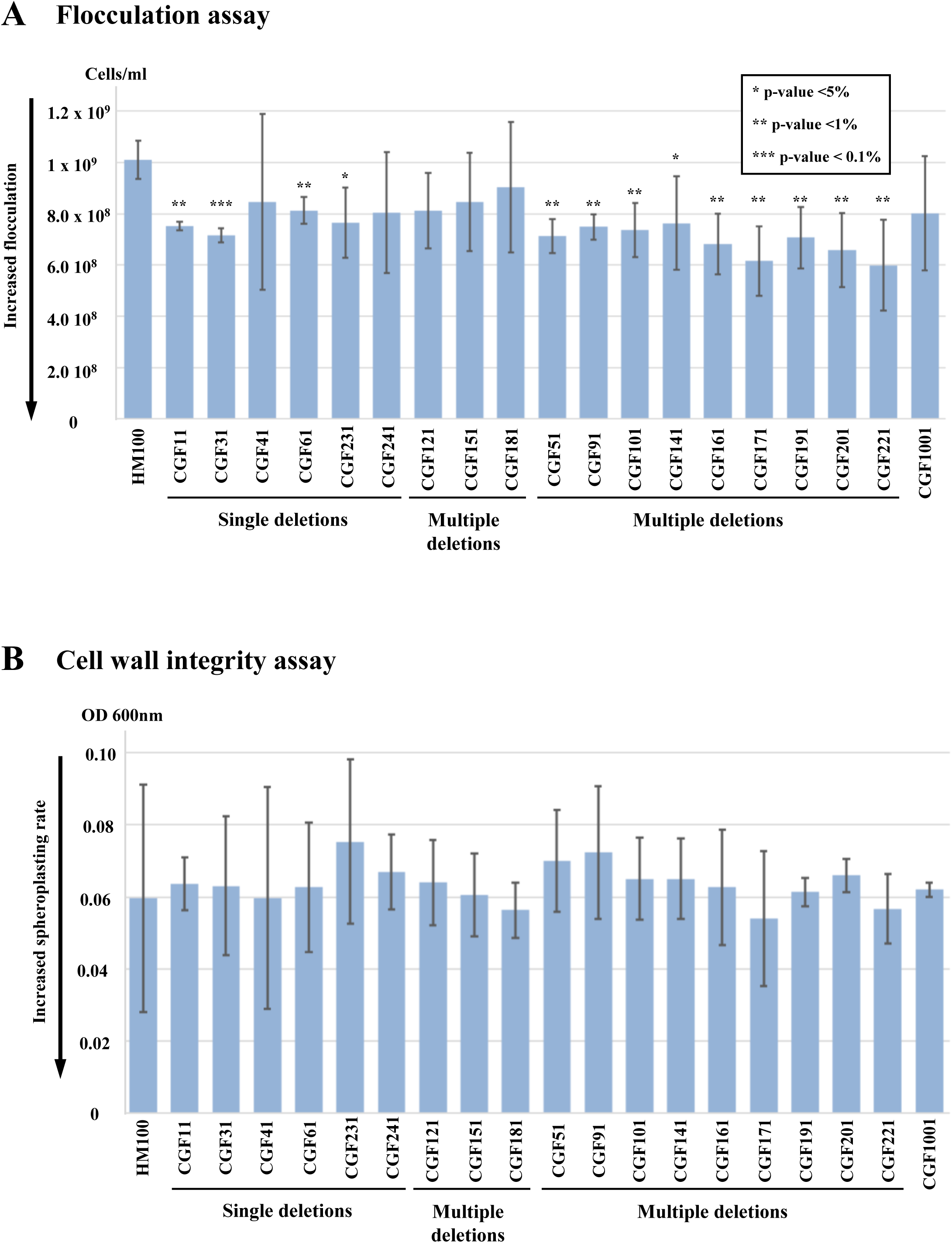
Flocculation and cell integrity experiments in *C. glabrata* mutants. **A**: Flocculation. Cell concentration as determined by optical density at 600 nm was determined for each strain. Error bars correspond to 95% confidence intervals. Student t-tests comparing each mutant to the reference HM100 strain were performed, asterisks showing significance levels: * p-value< 5%, ** p-value< 1%, *** p-value< 0.1%. **B**: Cell wall integrity. Optical density at 600 nm after zymolyase treatment is shown for each strain. Error bars are one standard deviation. No statistically significant difference was found among strains, by a Student t-test.

A spheroplast rate assay was performed in order to determine whether some mutations could significantly affect cell wall integrity. Optical densities at 600 nm was measured after incubation with a solution of zymolyase (Materials & Methods), OD reduction reflecting decrease in cell wall integrity (Ovalle *et al*. 1998). Overall, no significant difference was observed between any of the single or multiple mutants and the wild type control strain, or among the different mutants (Figure 4B). We concluded that none of the deleted genes was essential for cell wall integrity in *C. glabrata*.

### Megasatellite evolution of *S. cerevisiae* populations grown in bioreactor

In a second series of experiments, we decided to use *S. cerevisiae* as a test tube to address the intriguing question of megasatellite formation and expansion. Budding yeast encodes five genes involved in flocculation and cellular adhesion. *FLO1*, *FLO5* and *FLO9* are paralogues, each containing a 135 bp threonine-rich megasatellite (Richard and Dujon 2006). *FLO1* is the main gene responsible for flocculation and its efficacy is correlated to the megasatellite length (Verstrepen *et al*. 2005). *FLO10* encodes an 81 bp serine-rich minisatellite, whose sequence is unrelated to *FLO1*/5/9. *FLO11* encodes a highly repeated serine-rich 30 bp minisatellite, whose sequence is also unrelated to *FLO1*/5/9 but directly responsible for cellular adhesion of yeast cells with each other in specific conditions (Fidalgo *et al*. 2006). We wanted to delete all FLO megasatellites, in order to follow evolution of the sole *FLO1* gene in a bioreactor, over hundreds of generations. To that end, we engineered a wild type BY4741 strain as follows. First, we replaced the non-functional allele (*flo8-1*) in BY4741, with wild-type *FLO8*, which encodes *FLO1* transcriptional activator, in order to restore the strain capacity to flocculate (Liu *et al*. 1996). Next, *FLO9* and *FLO11* were deleted and the megasatellites encoded in *FLO5* and *FLO10* coding regions were perfectly deleted, leaving the remaining parts of both genes intact and in frame. The resulting strain (CSY2, Supplemental Table 3) flocculates very well in the presence of Ca^2+^. From this strain, a series of mutants were built, in which the *FLO1* megasatellite was replaced by one SHITT motif from *C. glabrata*, one ALS motif from *C. albicans*, one or two FLO motifs from *S. cerevisiae* or no motif at all (Figure 5A). These megasatellite motifs were chosen because they exhibit similar lengths (135 bp for FLO and SHITT, 108 bp for ALS), and because the FLO motif is the most widely spread in fungal genomes (Tekaia *et al*. 2013). None of these engineered strains was able to flocculate efficiently, proving that *FLO1* function depends on the FLO megasatellite for flocculation. These five strains, as well as a wild type BY4741 control and its *flo1*Δ derivative, were incubated in a bioreactor under constant oxygenation, in such a way that the air vent reached the bottom of the culture tube, in order to give a slight selective advantage to sedimenting cells (Figure 5B). Cultures were grown in parallel until flocculation was clearly visible as a drop in OD600 absorbance at the top of the tube, indicating that cell clumps were sedimenting to the bottom (Supplemental Figure 1). When a [Flo+] revertant appeared in one of the cultures, it was isolated and identified either by functional complementation with wild-type versions of genes known to inhibit flocculation, or by whole-genome sequencing if complementation did not suppress flocculation.

**Figure 5:**
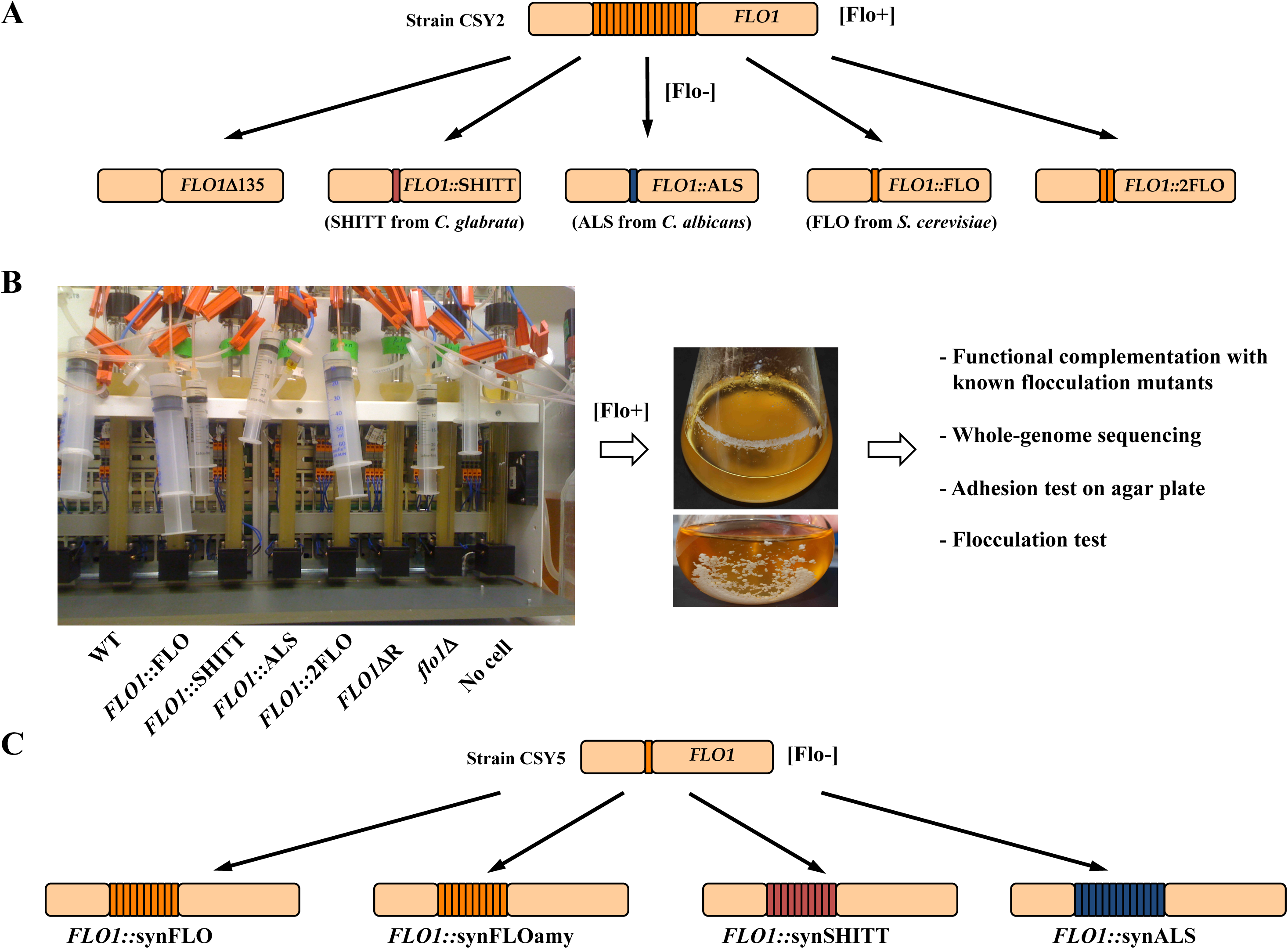
Evolution toward flocculation experimental setup. **A**: The wild-type *FLO1* gene was modified by the two-step replacement method, in such a way that the regular megasatellite was entirely deleted or replaced with one SHITT, one ALS, one FLO or two FLO motifs. Replacements did not disrupt the reading frame. **B**: Mutant strains as well as wild-type and *flo1*Δ controls were incubated in parallel bioreactors under constant oxygenation. The oxygen vents reached the bottom of each glass tube; in order to give a selective advantage to flocculating cells. Each time a [Flo+] mutant appeared, it was confirmed by a larger culture in flask allowing to assess the presence of large flocs each containing millions of yeast cells. These mutants were subsequently tested for dominance/recessivity and complementation tests with known flocculation mutants. **C**: Strain CSY5, containing a *FLO1* gene with only one FLO motif, was modified by the two-step replacement method so that the *FLO1* synthetic gene contained 10 copies in tandem of the FLO motif, of the SHITT motif or of the FLOamy motif, or 13 copies of the ALS motif, so that the total megasatellite length is approximately the same in all cases.

Three independent flocculation mutants were found in the BY4741 strain, after 48, 56 or 70 generations (Table 1): a point mutation in the *SSN6* gene, a general transcriptional corepressor (Chen *et al*. 2013), a short deletion in the *ACE2* gene, a transcription factor whose disruption prevents mother-daughter cell separation, generating multicellular yeast aggregates (Oud *et al*. 2013 p. 2; Ratcliff *et al*. 2015), and a point mutation in the *SRB8* gene, encoding a subunit of the RNA polymerase II mediator complex, involved in general transcriptional regulation, interacting with the Ssn6p-Tup1 complex (Núñez *et al*. 2007). This proved that our experimental setting was properly working to select flocculation proficient revertants. All strains were grown in parallel and an unexpected [Flo+] revertant was identified in the *flo1*Δ strain, after 70 generations. It turned out to be a point mutation in the *TUP1* gene, the *SSN6* partner in transcriptional repression. Another mutant arose in the *FLO1*::SHITT strain, after 218 generations. Whole-genome sequencing identified a 230 kilobases segmental duplication on chromosome II, extending from a Ty2 (*YBL100c*) to a Ty1 retrotransposon (*YBR013c*). Segmental duplications occurring between transposons or LTRs are frequent in *S. cerevisiae* (Koszul *et al*. 2004) and involve break-induced replication (Payen *et al*. 2008) but this one surprisingly covered the centromeric region, suggesting that the duplication was episomal, as was sometimes observed in evolution experiments with *S. cerevisiae* (Thierry *et al*. 2015). We did not investigate further this duplication. No other flocculation revertant could be identified in any of the other strains after more than 200 generations (Table 1), not even in the strain containing a tandem repeat of two FLO motifs in which we naively expected to detect an amplification by classical replication or recombination slippage (Richard and Pâques 2000). This result led us to the conclusion that generating a megasatellite by local duplication of a motif must be an extremely rare event, or that it occurred by a totally different mechanism than the one initially imagined, or only under particular environmental conditions.

### Functional variability of synthetic megasatellites in *S. cerevisiae*

In a third series of experiments, we determined whether a specific function could be attributed to a given megasatellite, or if any tandem repeat could be substituted while retaining the same general gene function. To that end, the *FLO1* gene was engineered to encode different synthetic megasatellites: 10 FLO repeats (hereafter called synFLO), 10 FLO repeats modified to encode amyloid-forming peptide motifs (synFLOamy) (Ramsook *et al*. 2010), 10 SHITT repeats (synSHITT) or 13 ALS repeats (synALS) (Supplemental Table 4). The *FLO1* gene was replaced in the CSY5 strain by each one of these four synthetic constructs in four different strains (Figure 5C). All these synthetic repeat strains, as well as the other strains used for evolution experiments were tested for four different phenotypes: adhesion to epithelial Lec2 cells, flocculation, invasion of agar plates and cell wall integrity (Figure 6).

**Figure 6:**
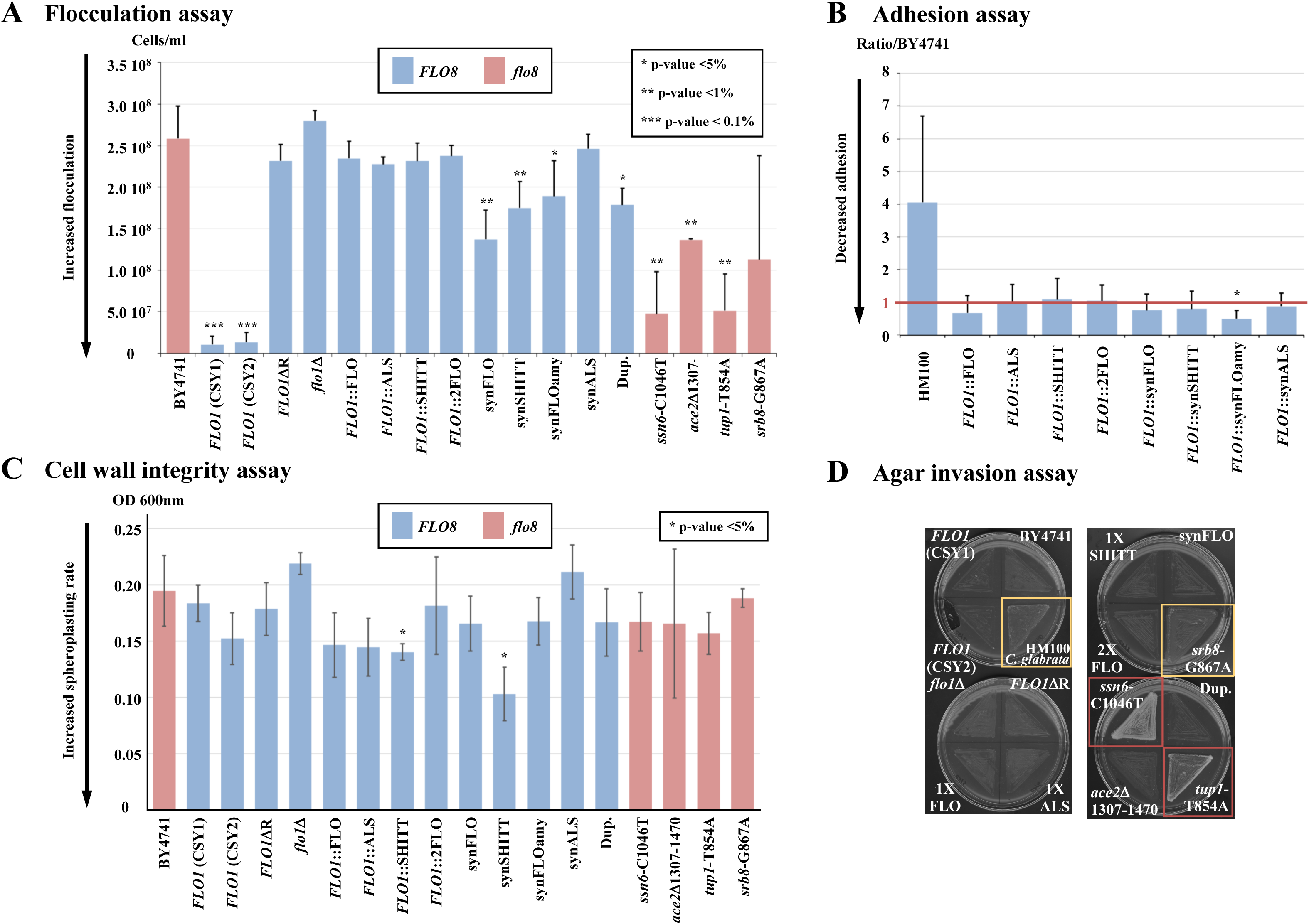
Results of phenotypic tests on mutant *FLO1* strains. **A**: Flocculation. Cell concentration as determined by optical density at 600 nm was determined for each strain. Error bars correspond to 95% confidence intervals. Student t-tests comparing each mutant to the reference BY4741 strain were performed, asterisks showing significance levels: * p-value< 5%, ** p-value< 1%, *** p-value< 0.1%. **B**: Adhesion to epithelial cells. Experiments were performed as in Figure 3, and results are shown as ratios to BY4741 reference strain. Error bars correspond to 95% confidence intervals. Non-parametric Mann-Whitney tests were performed, the only strain statistically different from BY4741 contains the synthetic FLOamy construct and is marked by an asterisk (p-value <5%). **C**: Cell wall integrity. Optical density at 600 nm after zymolyase treatment is shown for each strain. Error bars are one standard deviation. Student t-tests comparing each mutant to the reference BY4741 strain were performed, asterisks showing significant differences (p-value < 5%). **D**: Invasion of agar plates. Pictures of each plate were taken 24 hours after water washes (Materials & Methods). Two strains show weak adhesion (HM100 and *srb8*-G867A, yellow squares) and two strains show strong adhesion (*ssn6*-C1046T and *flo1*Δ *tup1*-T854A, red squares).

Results obtained for flocculation showed that none of the *FLO1* gene with only one motif (FLO, ALS, SHITT) or two motifs (2FLO) flocculated better than a strain in which all *FLO1* repeats were deleted (*FLO1*ΔR strain, Figure 6A). [Flo+] revertants that were isolated during the previous evolution experiment (strains CSY20 to 24) flocculated well, although at variable levels and always less than *FLO8 FLO1* cells (CSY1 or CSY2). Interestingly, synthetic repeat strains displayed different flocculation degrees; synFLO, synSHITT and synFLOamy were statistically different from wild type, but synALS did not flocculate, proving that although its total length was similar to the three others this megasatellite is not sufficient to trigger flocculation. Therefore, this phenotype was dependent on the particular sequence of the megasatellite used rather than on a fixed distance between N- and C-terminus of the encoded protein.

Next, we investigated the adhesion to epithelial cells of the different engineered *FLO1* mutants. As expected, *C. glabrata* (HM100) was more adherent than any of the *S. cerevisiae* strains (Figure 6B). None of the FLO, SHITT or ALS strains increased adherence over background. The synFLOamy construct, but not the synFLO construct, exhibited a significant decrease in adhesion, as compared to wild type, showing that in the context of *FLO1*, the corresponding megasatellite partially inhibited adhesion to epithelial cells. It is possible that facilitating yeast-yeast interactions by making amyloid fibers decreased possible interactions between yeast and epithelial cells.

Cell wall integrity was assayed by zymolyase-induced spheroplast rate, as previously. Only two strains were statistically different from wild type, CSY7 and CSY10, encoding respectively *FLO1*::SHITT and *FLO1*::synSHITT motifs. We concluded that the SHITT motif from *C. glabrata* modified Flo1p function in such a way that budding yeast cell wall integrity was slightly altered (Figure 6C).

Finally, invasion of agar plates was also tested. *C. glabrata* (HM100) and the *srb8*-G867A mutant were moderately invasive, whereas *ssn6*-C1046T and *tup1*-T854A mutants were more invasive than any other strain (Figure 6D). We concluded that these three mutations that increased flocculation also increased agar invasion, whereas none of the synthetic strains (in which *FLO11* and *FLO9* were deleted) showed any invasion of agar plates.

## DISCUSSION

### Subtelomeric megasatellites of the same paralogous family exhibit opposite roles in adhesion to epithelial cells

Previous experiments showed that in log phase cells, at least in some strains, *EPA1* encodes the main adhesin in *C. glabrata* (Cormack *et al*. 1999). Two other subtelomeric genes, *EPA6* and *EPA7*, are involved in adherence, these genes being normally repressed by subtelomeric silencing involving the *SIR3* and *RIF1* genes (Castano *et al*. 2005), as well as a negative regulator element located at 3’ ends of these genes (Gallegos-García *et al*. 2012). Subsequent structure-function studies of *EPA1* showed that the length of the Ser/Thr rich region tandemly repeated in the middle of the protein was important for cellular adhesion, since when it was shortened below 100 amino acids adhesion was lost. This was interpreted by the need for the ligand binding N-terminal domain to be projected away from the membrane-attached domain, in order to interact with the extracellular environment (Frieman *et al*. 2002). In our present experiments, we tested whether deleting genes with longer megasatellites would have a more drastic impact on adhesion but no correlation was found between length of megasatellite in a deleted gene and impact on adhesion to epithelial cells (Supplemental Figure 2). In addition, several mutants showed an increased adhesion as compared to wild type (Figure 3B). This was the case for genes *CAGL0B05061g*, *CAGL0K13024g*, and *CAGL0A04851g*. Note that *CAGL0K13024g* deletion was shown here to increase adhesion, while mass spectrometry analysis of *C. glabrata* cell wall peptides identified its product as a *bona fide* cell wall component (Kraneveld *et al*. 2011). A similar observation was made when deletion of the *FIG2* gene increased cellular adhesion in *S. cerevisiae* (Jue and Lipke 2002). The authors concluded that the glycosylated part of the Fig2 protein extended far from the cell surface, “hiding” residues of proteins involved in adhesion. Hence, removing Fig2 would unmask these residues, thus increasing adhesion.

It was previously reported that overexpressing *FLO1* or *FLO5* in *S. cerevisiae* may lead to opposite effects in coaggregation experiments with other yeasts, such as *Lachancea thermotolerans* and *Hanseniaspora opuntiae*. This led to the conclusion that paralogous flocculins exhibit different properties in complex ecosystems containing more than one yeast species (Rossouw *et al*.). In our experiments, all the megasatellite-containing genes studied are paralogues (except *CAGL0C00253g* and *CAGL0K13024g*, Figure 1C). The N-terminal and C-terminal parts of their encoded proteins are therefore homologous (Rolland *et al*. 2010). However, some deletions are associated to increased adhesion while others have the opposite effect. This suggests that these paralogues play different roles in the pathogenic life of *C. glabrata*, faced with distinctive challenges when infecting a human host.

It was previously shown that different *C. glabrata* isolates displayed a wide range of adhesion properties in a mouse model of infection (Atanasova *et al*. 2013), which is not surprising given the high genomic variability of this clade (Muller *et al*. 2009; Gabaldon *et al*. 2013; Carreté *et al*. 2019). However, in our present experiments, all strains were built from the CBS138 reference strain and are therefore isogenic, except for the subset of deleted megasatellite-containing genes. It is therefore striking that deletion of multiple megasatellite containing genes did not give a cumulative phenotype. In particular, the strain in which 11 megasatellite-containing genes were deleted (CGF221) did not show any adherence defect as compared to wild type.

Each of these megasatellite-encoded proteins may be present in variable amounts at the cell surface, and their role in adhesion may depend in complex ways on the total complement of cell wall proteins. It is possible that those not directly involved in adhesion may alter the cell wall surface and modify its properties. However, zymolyase experiments show that none of these deletions induces a detectable decrease in cell wall integrity, even the large 11-gene deletion. Therefore, it is possible that this category of genes plays a more important role in different physiological conditions or in more complex ecological niches. It is interesting to note that adhesion to epithelial cells or to other *C. glabrata* cells are not mediated by the same genes to the exception of *CAGL0B05061g* and *CAGL0A04851g*, suggesting that these two genes negatively control all cell-to-cell interactions.

### Directed evolution experiments do not select megasatellite amplification

Former evolution experiments in *S. cerevisiae* used either limiting growth condition, like glucose (Dunham *et al*. 2002), gene dosage assay of a ribosomal protein (Koszul *et al*. 2004) or partially deficient tRNA synthetase (Thierry *et al*. 2015) to select mutants that would grow like wild type in challenging conditions. Chromosomal duplications of large DNA segments were frequently observed, involved retrotransposons, LTRs or microsatellites (Payen *et al*. 2008). In the present evolution experiments, we expected to select the local amplification of a megasatellite seed inserted in the *FLO1* gene, by selecting [Flo+] revertants from [Flo-] cells. *FLO1* was chosen because it shows a wide variety of phenotypes directly correlated to expression level (Smukalla *et al*. 2008) and to megasatellite length (Verstrepen *et al*. 2005). [Flo+] revertants involving megasatellite amplification were never observed after more than 200 generations. Instead, revertants corresponded to mutations in general transcription factors such as *SSN6*-*TUP1* (Chen *et al*. 2013), *SRB8* (Núñez *et al*. 2007) or *ACE2* (Ratcliff *et al*. 2012, 2015; Oud *et al*. 2013). All these mutations happened in the control BY4741 or its *flo1*Δ derivative, mutated for the *FLO8* transcription activator (Table 1 and Supplemental Table 3). The only [Flo+] revertant that was identified in one of the non-control strains was a segmental duplication of 230 kb on chromosome II, encompassing more than 100 genes and involving two Ty elements, reminiscent of similar duplications in other experimental systems (Dunham *et al*. 2002; Koszul *et al*. 2004; Thierry *et al*. 2015). No local duplication of a megasatellite seed was detected, not even in the strain containing two FLO motifs that could easily duplicate by replication or recombination slippage (Richard and Pâques 2000; Richard *et al*. 2008). Note that in our experiments, there is no limiting factor, cells were grown in rich medium under constant oxygenation. These conditions of rapid growth may favor large segmental duplications over local slippages. Growing cells in more stressful conditions or at a lower temperature to slow down replication may increase chances to detect other kinds of mutations.

### Differential functions of synthetic megasatellites in *S. cerevisiae*

One striking result of our experiments is that one given megasatellite may not be replaced by another one of the same length without losing some cell properties. The fact that the synFLO, synSHITT and synFLOamy strains all flocculated while synALS did not flocculate, shows that one tandem repeat may not necessarily substitute to another one of the same length to perform the same function (Figure 6A). Similarly, the synFLOamy strain exhibited reduced adhesion to epithelial cells as compared to the synFLO strain and other synthetic strains. This shows that replacing the FLO megasatellite by megasatellites from adherent pathogenic yeasts (synALS and synSHITT) is not sufficient to increase *S. cerevisiae* adhesion to epithelial cells, unlike expressing intact *EPA1*, for example (Cormack *et al*. 1999). These data demonstrate that the tested megasatellites themselves are not able to increase adherence to epithelial cells, but almost certainly work in conjunction with other domains in their resident protein to carry out this function.

In previous work comparing different phenotypes of natural *S. cerevisiae* isolates, the authors also concluded that no correlation could be found between flocculation and invasion phenotypes (Hope and Dunham 2014). However, large genetic differences existed between the different isolates. In our experiments all strains are perfectly isogenic except for the *FLO1* megasatellite sequence. We also found no correlation between the ability to flocculate (synFLO, synSHITT and synFLOamy all flocculate) and to invade agar (none of the synthetic strains were able to invade).

### Conclusions

In all, our study shows that megasatellites contribute to cell surface phenomena like adherence and flocculation, but in a complex manner. While, for example, we could document a role for particular megasatellite genes in adherence, the deletion of 11 megasatellite genes in CBS138 did not strongly alter its adherence, nor its cell wall integrity. We also found a role for megasatellite repeats in function of the flocculin Flo1, and show that function was affected by particular megasatellite sequences, as opposed to these sequences simply acting as a spacer of a given length. Lastly, we did not find evidence that any of the megasatellite repeats tested were direct mediators of adherence, or agar invasion, since their expression in the context of the Flo1 flocculin permitted flocculation but not adherence or agar invasion. It seems likely, therefore, that they contribute by functioning with other domains of the proteins in which they are encoded.

## Supporting information

Table 1. List of mutants selected.

## ACKNOWLEDGEMENTS

This work was supported by the Institut Pasteur, the Centre National de la Recherche Scientifique (CNRS), by an ANR grant to B.D. (project DYGEVO) and by grant R01AI046223 to B.P.C. We wish to thank Maria Teresa Teixeira for helpful advices to design the telomeric disruption plasmid, pMEG3.

