## Supplementary material for "Functional variability in adhesion and flocculation of yeast megasatellite genes": Table 1. List of mutants selected.

**Table 1: List of [Flo+] revertants identified during evolution of *S. cerevisiae* strains**

| **FLO1 allele** | **Nbr generations** | **Mutants** | **Strain name** |
| --- | --- | --- | --- |
| WT (BY4741) | 56 | *ssn6*-C1046T | CSY20 |
|  | 48 | *ace2*Δ(1307-1470) | CSY21 |
|  | 70 | *srb8*-G867A | CSY24 |
| *FLO1*::FLO | 214 | - | - |
| *FLO1*::SHITT | 218 | YBL100c-YBR13c dup (230 kb) | CSY23 |
| *FLO1*::ALS | 216 | - | - |
| *FLO1*::2FLO | 217 | - | - |
| *FLO1*::Δ135 | 214 | - | - |
| *flo1*Δ | 70 | *tup1*-T854A | CSY22 |
