## Supplemental Figure 1 for "Functional variability in adhesion and flocculation of yeast megasatellite genes"

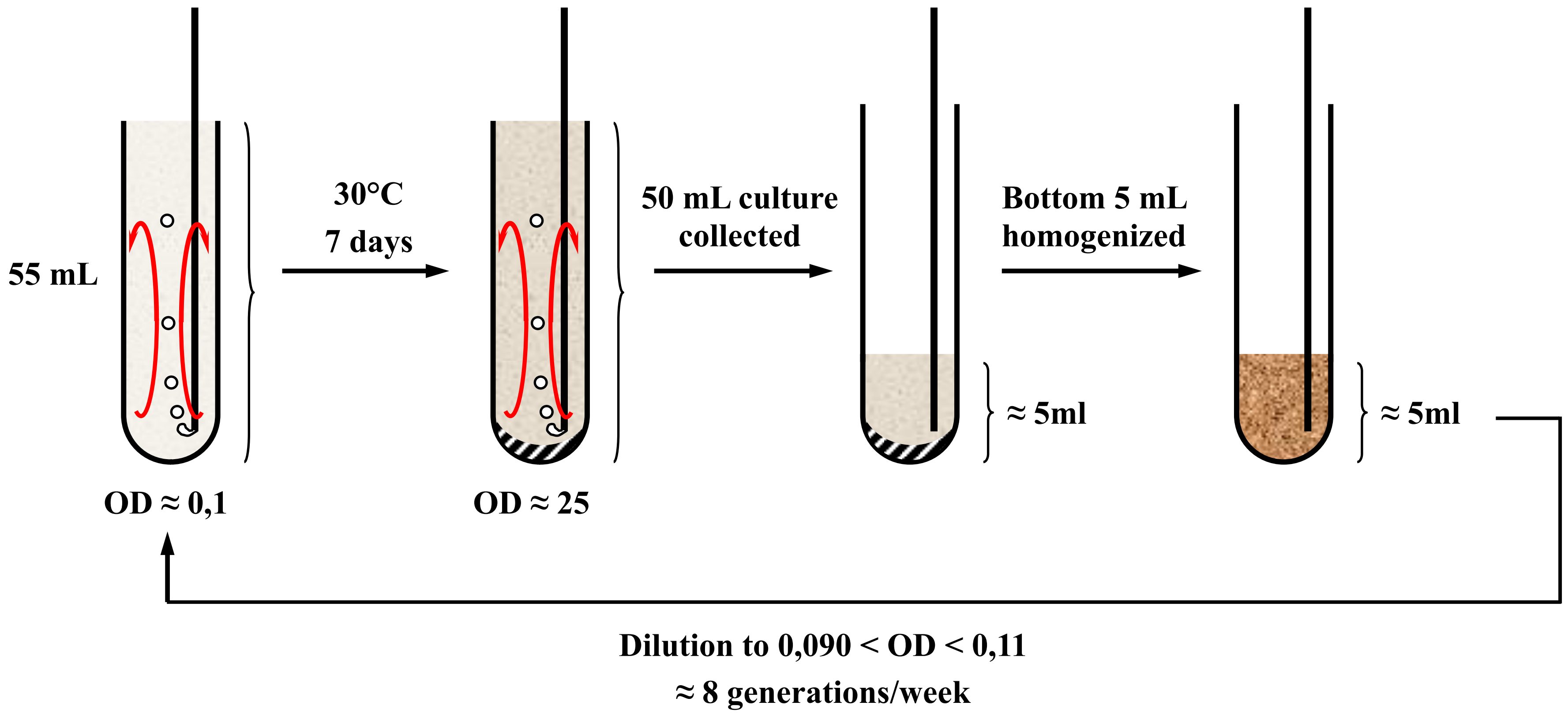


**Supplemental Figure 1: Protocol of *in vitro* evolution of *S. cerevisiae* mutants**

Yeast cells were grown in 55 mL bioreactors, in YPD. Stationary phase was reached around OD 25 (1.8 x $10^8$ cells/mL). After one week, 50 mL of culture was removed from the top of the bioreactor. The bottom 5 mL was homogenized and diluted to OD ~0.1 (7 x $10^5$ cells/mL) in fresh YPD.
