## Supplemental Figure 2 for "Functional variability in adhesion and flocculation of yeast megasatellite genes"

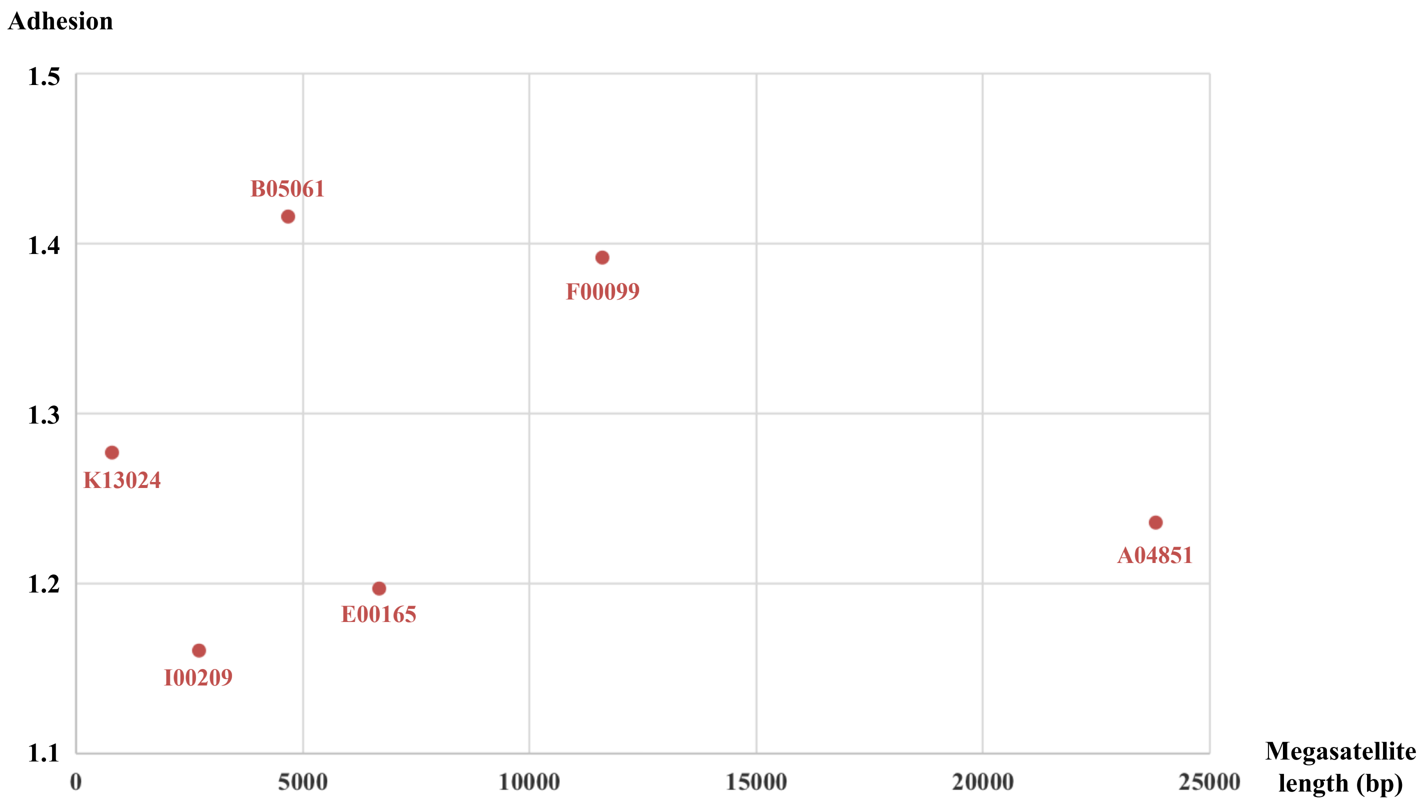


**Supplemental Figure 2: Absence of correlation between adhesion and megasatellite length**
Adhesion of each mutant strain as compared to wild-type control (Y axis) was plotted as a function of megasatellite length (X axis). Genes are shown in their abbreviated form (see Figure 3).
