## Supplemental Table 1 for "Functional variability in adhesion and flocculation of yeast megasatellite genes"

**Supplemental Table 1: Sequence of PCR primers used in this study and corresponding plasmids**

| **Name** | **DNA sequence** | **Plasmid** |
| --- | --- | --- |
| YALIupfor | GGGGTACCCCCAGCAGCGTGTCTATGGGTTCT | pMEG0 |
| YALIuprev | CGGGATCCCGCGTTGGCCGCCCAAACGTCTCC |  |
| CgURA3for | GGAAGATCTTCCTCCTGTAATTACAACAATTCAA | pMEG1 |
| CgURA3rev | CGGGATCCCGGTTGCCATTACGCCACGCGAGC |  |
| YALIdownfor | GGAAGATCTTCCCAGCAGCGTGTCTATGGGTTCT | pMEG2 |
| YALIdownrev | ATAAGAATGCGGCCGCTAAACTATCGTTGGCCGCCCAAACGTCTCC |  |
| TELup | GGCCTGGGGTCTGGGTGCTGTGGGGTCTGGGTGCTGTGGGGTCTGGGTGCTGTGGGGTCTGGGTGCTGTGGGGTCTGGGTGCTG | pMEG3 |
| TELdown | GGCCCAGCACCCAGACCCCACAGCACCCAGACCCCACAGCACCCAGACCCCACAGCACCCAGACCCCACAGCACCCAGACCCCA |  |
| **Name** | **DNA sequence** |  |
| A04851for | GGGGTACCGCGGatgagattcaaaaggatacttt | pMEG12 |
| A04851rev | GGGGTACCTATGAAGTCGAGTTGCAATGAT |  |
| B05061for | GGGGTACCGCGGATGAAGTACAAAACCCACCCTC | pMEG4 |
| B05061rev | GGGGTACCGACTATTTGTTAAAGTGGTATA |  |
| E00165for | GGGGTACCGCGGATGCTATTGCGAAATATTTACC | pMEG13 |
| E00165rev | GCCGGTACCCCATATTCATTAGTATA |  |
| F00099for | GGGGTACCGCGGATGATGAGAAAAAAGCCACCAT | pMEG5 |
| F00099rev | GGGGTACCGTCAAACCCAGTCACCCTGCTA |  |
| G10219for | GGGGTACCGCGGAGTTCAACTATCATGGTGGTTC | pMEG15 |
| G10219rev | GGGGTACCTATCATCGTTTCTTCGATGTAT |  |
| H00209for | GGGGTACCGCGGATCACAAAACAACGAAGAAATA | pMEG6 |
| H00209rev | GGGGTACCTTTAGAATAAGATATA |  |
| H10626for | GGGGTACCGCGGATGAGATTTAAAAGTATATTTT | pMEG16 |
| H10626rev | GGGGTACCTGGTGGGGCGTAGAGGTAGACT |  |
| I00209for | GGGGTACCGCGGCTTCGCAACAAATGGCCTCCAT | pMEG11 |
| I00209rev | ATTGATAGAGGTACCTAAACCAGTA |  |
| L00157for | GGGGTACCGCGGATGAGATTTAGAAACATATTAT | pMEG17 |
| L00157rev | GGGGTACCAAATTCCGATTATACGACGACG |  |
| L00227for | GGGGTACCGCGGTCTTGAATGACATTTACTAAGA | pMEG18 |
| L00227rev | GGGGTACCCAAAGAATAGATAAGACCGGTA |  |
| K13024for | GGGGTACCGCGGATGAGGCTTTATCGGTGTTTTT | pMEG9 |
| K13024rev | GGGGTACCCGGAATTTTTCTGCTGTTTTGT |  |
| EPA1for | GGGGTACCGCGGCACGGAGAAGTTTTTCTGTGGA |  |
| EPA1rev | GGGGTACCTGTTTTAGTTTGGTTAATTGCA |  |

**Top:**

YALIup primers: underlined sequences correspond to KpnI and BamHI cloning sites (see Materials & Methods).

CgURA3 primers: underlined sequences correspond to BglII and BamHI cloning sites.

YALIdown primers: underlined sequences correspond to BglII and NotI cloning sites.

TEL primers: underlined sequences indicate telomeric repeat borders.

**Bottom:**

Megasatellite deletion primers: underlined sequences indicate KpnI sites used for cloning into pMEG3.
