## Supplemental Table 2 for "Functional variability in adhesion and flocculation of yeast megasatellite genes"

**Supplemental Table 2: Sequence of PCR primers used in this study to build synthetic *FLO1* plasmids**

| **Name** | **DNA sequence** |
| --- | --- |
| SC1 | atacgactcactatagggcgaattgggtaccGGGCCCcccCTCGAGCATGCtcactattaggcttgttaaag |
| SC2 | tggTTCCGTTGTAGTTGTAG |
| SC3 | TCAGGACAAATCACCAGCTC |
| SC4 | accctcactaaagggaacaaaagctgGAGCTCcaccgcggTCTAGAAGCTtaagtactcatacatcggtttc |
| SC5 | CCACTACAACTACAACGGAACCATCAGGACAAATCACCAGCTC |
| SC6 | GAGCTGGTGATTTGTCCTGATGGTTCCGTTGTAGTTGTAGTGG |
| SC7 | AAATTATGCTGTCAGTACCACTACAACTACAACGGAACCATGGACTGGTACTGAAACCTC |
| SC8 | GGTAGTTGGCTTACCGTCAGAACCGGTAAAGGTGGTAGTTTCAGTAGAAAATGTAGAGGTTTCAGTACCAGTCCA |
| SC9 | TGACGGTAAGCCAACTACCGAAACCATCTACTACGTCGAGACACCAACAGTCGGTACTGCCTCTACTACCTACAC |
| SC10 | TAATTGGACGCGAAGACGTGATAGAGCTGGTGATTTGTCCTGATGGAGTGTAGGTAGTAGAGGCAGT |
| SC11 | AAATTATGCTGTCAGTACCACTACAACTACAACGGAACCAGCCGGTGAAGCCGACTACACTACTACC |
| SC12 | GTGGTAGTGATGTGAGAGACCAAATCGGTTTCAAAGTCACCGTTACCCTTAGTGATGGTAGTAGTGTAGTCGGCT |
| SC13 | GTCTCTCACATCACTACCACAGACAGCGATGGTAAGCCAACTACCATTACTACCACAATTCCATTGGACGAC |
| SC14 | TAATTGGACGCGAAGACGTGATAGAGCTGGTGATTTGTCCTGAGTCGTCCAATGGAATTGTGG |
| SC15 | AAATTATGCTGTCAGTACCACTACAACTACAACGGAACCAAGAGAACCACCAAACCCAAC |
| SC16 | TGTGGCGTAAGACTGAGACCAGTATTCGGTGGTAGTGACAGTTGGGTTTGGTGGTTCTCT |
| SC17 | GGTCTCAGTCTTACGCCACAACTACTACCGTCACCGCCCCACCAGGTGGTACTGACACAG |
| SC18 | TAATTGGACGCGAAGACGTGATAGAGCTGGTGATTTGTCCTGAGATGATAACTGTGTCAGTACCACCTGG |
| SC19 | TGGACTGGTACTGAAACCTC |
| SC20 | TGGAGTGTAGGTAGTAGAGGCAGT |
