## Supplemental Table 3 for "Functional variability in adhesion and flocculation of yeast megasatellite genes"

**Supplemental Table 3: List of strains used in this study**

| **Species** | **Strain** | **Genotype** | **Origin** |
| --- | --- | --- | --- |
| *C. glabrata* | CBS138 | Wild type | Collection* |
| *C. glabrata* | HM100 | *ura3*Δ::KANMX | (Muller et al., 2008) |
| *C. glabrata* | BG2 | Wild type | (Cormack et al., 1999) |
| *C. glabrata* | BG14 | *ura3*Δ (-85 + 932)::Tn903NeoR | BG2 |
| *C. glabrata* | BG64 | *ura3*Δ (-85 + 932)::Tn903NeoR *epa1*Δ | BG2 |
| *C. glabrata* | BG923 | *ura3*Δ (-85 + 932)::Tn903NeoR *epa1*Δ *epa6*Δ *epa7*Δ | (Castano et al., 2005) |
| *C. glabrata* | CGF11 | *ura3*Δ::KANMX *CAGL0B05061*Δ | HM100 |
| *C. glabrata* | CGF21 | *ura3*Δ (-85 + 932)::Tn903NeoR *CAGL0B05061*Δ | BG2 |
| *C. glabrata* | CGF31 | *ura3*Δ::KANMX *CAGL0F00099*Δ | HM100 |
| *C. glabrata* | CGF41 | *ura3*Δ::KANMX *CAGL0K13024*Δ | HM100 |
| *C. glabrata* | CGF51 | *ura3*Δ::KANMX *CAGL0B05061*Δ *CAGL0K13024*Δ | CGF11 |
| *C. glabrata* | CGF61 | *ura3*Δ::KANMX *CAGL0A04851*Δ | HM100 |
| *C. glabrata* | CGF91 | *ura3*Δ::KANMX *CAGL0B05061*Δ *CAGL0K13024*Δ *CAGL0H00209*Δ | CGF51 |
| *C. glabrata* | CGF101 | *ura3*Δ::KANMX *CAGL0B05061*Δ *CAGL0K13024*Δ *CAGL0H00209*Δ *CAGL0G10219*Δ | CGF91 |
| *C. glabrata* | CGF121 | *ura3*Δ::KANMX *CAGL0A04851*Δ *CAGL0I00209*Δ | CGF61 |
| *C. glabrata* | CGF141 | *ura3*Δ::KANMX *CAGL0B05061*Δ *CAGL0K13024*Δ *CAGL0H00209*Δ *CAGL0G10219*Δ *CAGL0F00099*Δ | CGF101 |
| *C. glabrata* | CGF151 | *ura3*Δ::KANMX *CAGL0A04851*Δ *CAGL0I00209*Δ *CAGL0E00165*Δ | CGF121 |
| *C. glabrata* | CGF161 | *ura3*Δ::KANMX *CAGL0B05061*Δ *CAGL0K13024*Δ *CAGL0H00209*Δ *CAGL0G10219*Δ *CAGL0F00099*Δ *CAGL0H10626*Δ | CGF141 |
| *C. glabrata* | CGF171 | *ura3*Δ::KANMX *CAGL0B05061*Δ *CAGL0K13024*Δ *CAGL0H00209*Δ *CAGL0G10219*Δ *CAGL0F00099*Δ *CAGL0H10626*Δ *CAGL0L00227*Δ *CAGL0L00157*Δ | CGF161 |
| *C. glabrata* | CGF181 | *ura3*Δ::KANMX *CAGL0A04851*Δ *CAGL0I00209*Δ *CAGL0E00165*Δ *CAGL0L00157*Δ | CGF151 |
| *C. glabrata* | CGF191 | *ura3*Δ::KANMX *CAGL0B05061*Δ *CAGL0K13024*Δ *CAGL0H00209*Δ *CAGL0G10219*Δ *CAGL0F00099*Δ *CAGL0H10626*Δ *CAGL0L00227*Δ *CAGL0L00157*Δ *CAGL0A04851*Δ | CGF171 |
| *C. glabrata* | CGF201 | *ura3*Δ::KANMX *CAGL0B05061*Δ *CAGL0K13024*Δ *CAGL0H00209*Δ *CAGL0G10219*Δ *CAGL0F00099*Δ *CAGL0H10626*Δ *CAGL0L00227*Δ *CAGL0L00157*Δ *CAGL0A04851*Δ *CAGL0I00209*Δ | CGF191 |
| *C. glabrata* | CGF221 | *ura3*Δ::KANMX *CAGL0B05061*Δ *CAGL0K13024*Δ *CAGL0H00209*Δ *CAGL0G10219*Δ *CAGL0F00099*Δ *CAGL0H10626*Δ *CAGL0L00227*Δ *CAGL0L00157*Δ *CAGL0A04851*Δ *CAGL0I00209*Δ *CAGL0E00165*Δ | CGF201 |
| *C. glabrata* | CGF231 | *ura3*Δ::KANMX *CAGL0E00165*Δ | HM100 |
| *C. glabrata* | CGF241 | *ura3*Δ::KANMX *CAGL0I00209*Δ | HM100 |
| *C. glabrata* | CGF251 | *ura3*Δ::KANMX *CAGL0L00157*Δ | HM100 |
| *C. glabrata* | CGF1001 | *ura3*Δ::KANMX *epa1*Δ0 *epa2*Δ0 *epa3*Δ0 | HM100 |
| *S. cerevisiae* | BY4741 | *ura3*Δ0 *his3*Δ1 *leu2*Δ0 *met15*Δ0 *flo8*-1 | (Brachmann et al., 1998) |
| *S. cerevisiae* | BY6870 | *ura3*Δ0 *his3*Δ1 *leu2*Δ0 *met15*Δ0 *flo8*-1 *flo1*ΔKANMX | (Winzeler et al., 1999) |
| *S. cerevisiae* | CSY1 | *ura3*Δ0 *his3*Δ1 *leu2*Δ0 *met15*Δ0 *FLO8* *flo9*Δ0 *flo11*Δ::KANMX *FLO5*ΔR | BY4741 |
| *S. cerevisiae* | CSY2 | *ura3*Δ0 *his3*Δ1 *leu2*Δ0 *met15*Δ0 *FLO8* *flo9*Δ0 *flo11*Δ::KANMX *FLO5*ΔR *FLO10*ΔR | CSY1 |
| *S. cerevisiae* | CSY3 | *ura3*Δ0 *his3*Δ1 *leu2*Δ0 *met15*Δ0 *FLO8* *flo9*Δ0 *flo11*Δ::KANMX *FLO5*ΔR *FLO10*ΔR *FLO1*ΔR | CSY2 |
| *S. cerevisiae* | CSY4 | *ura3*Δ0 *his3*Δ1 *leu2*Δ0 *met15*Δ0 *FLO8* *flo9*Δ0 *flo11*Δ::KANMX *FLO5*ΔR *FLO10*ΔR *flo1*Δ::*MET15* | CSY2 |
| *S. cerevisiae* | CSY5 | *ura3*Δ0 *his3*Δ1 *leu2*Δ0 *met15*Δ0 *FLO8* *flo9*Δ0 *flo11*Δ::KANMX *FLO5*ΔR *FLO10*ΔR *FLO1*::FLO | CSY2 |
| *S. cerevisiae* | CSY6 | *ura3*Δ0 *his3*Δ1 *leu2*Δ0 *met15*Δ0 *FLO8* *flo9*Δ0 *flo11*Δ::KANMX *FLO5*ΔR *FLO10*ΔR *FLO1*::ALS | CSY2 |
| *S. cerevisiae* | CSY7 | *ura3*Δ0 *his3*Δ1 *leu2*Δ0 *met15*Δ0 *FLO8* *flo9*Δ0 *flo11*Δ::KANMX *FLO5*ΔR *FLO10*ΔR *FLO1*::SHITT | CSY2 |
| *S. cerevisiae* | CSY8 | *ura3*Δ0 *his3*Δ1 *leu2*Δ0 *met15*Δ0 *FLO8* *flo9*Δ0 *flo11*Δ::KANMX *FLO5*ΔR *FLO10*ΔR *FLO1*::2FLO | CSY2 |
| *S. cerevisiae* | CSY9 | *ura3*Δ0 *his3*Δ1 *leu2*Δ0 *met15*Δ0 *FLO8* *flo9*Δ0 *flo11*Δ::KANMX *FLO5*ΔR *FLO10*ΔR *FLO1*::synFLO | CSY2 |
| *S. cerevisiae* | CSY10 | *ura3*Δ0 *his3*Δ1 *leu2*Δ0 *met15*Δ0 *FLO8* *flo9*Δ0 *flo11*Δ::KANMX *FLO5*ΔR *FLO10*ΔR *FLO1*::synSHITT | CSY2 |
| *S. cerevisiae* | CSY11 | *ura3*Δ0 *his3*Δ1 *leu2*Δ0 *met15*Δ0 *FLO8* *flo9*Δ0 *flo11*Δ::KANMX *FLO5*ΔR *FLO10*ΔR *FLO1*::synFLOamy | CSY2 |
| *S. cerevisiae* | CSY12 | *ura3*Δ0 *his3*Δ1 *leu2*Δ0 *met15*Δ0 *FLO8* *flo9*Δ0 *flo11*Δ::KANMX *FLO5*ΔR *FLO10*ΔR *FLO1*::synALS | CSY2 |
| *S. cerevisiae* | CSY20 | *ura3*Δ0 *his3*Δ1 *leu2*Δ0 *met15*Δ0 *flo8*-1 *ssn6*-C1046T | BY4741 |
| *S. cerevisiae* | CSY21 | *ura3*Δ0 *his3*Δ1 *leu2*Δ0 *met15*Δ0 *flo8*-1 *ace2*Δ(1307-1470) | BY4741 |
| *S. cerevisiae* | CSY22 | *ura3*Δ0 *his3*Δ1 *leu2*Δ0 *met15*Δ0 *flo8*-1 *flo1*ΔKANMX *tup1*-T854A | BY6870 |
| *S. cerevisiae* | CSY23 | *ura3*Δ0 *his3*Δ1 *leu2*Δ0 *met15*Δ0 *flo8*-1 *FLO8* *flo9*Δ0 *flo11*Δ::KANMX *FLO5*ΔR *FLO10*ΔR *FLO1*::SHITT *YBL100c*-*YBR013c* dup | CSY7 |
| *S. cerevisiae* | CSY24 | *ura3*Δ0 *his3*Δ1 *leu2*Δ0 *met15*Δ0 *flo8*-1 *srb8*-G867A | BY4741 |

* Collection: CBS-KNAW collection

All *S. cerevisiae* strains are *MAT*a

For *C. glabrata* strains, all CBS138 derivatives are *MAT*α and all BG2 derivatives are M*AT*a.

Note that all *C. glabrata* strains whose name ends by '1' are the [Ura-] product of the same [Ura+] mutant before 5-FOA selection, e. g. CGF11 comes from CGF1 whose genotype is *CAGL0B05061*Δ*URA3*. After 5-FOA selection, the *URA3* marker was lost and the resulting strain had a '1' added to its name to differentiate it from its mother. Only strains which lost the *URA3* marker were used in the present study and only these are shown in the above table.
