## Supplemental Table 4 for "Functional variability in adhesion and flocculation of yeast megasatellite genes"

**Supplemental Table 4: Sequence of megasatellite motifs used in this study**

| **Megasatellite motif** | **DNA sequence** | **Peptide sequence** |
| --- | --- | --- |
| FLO | TGGACCGGTACAGAGACTAGTACCTTTTCTACTGAGACAACTACCTTCACAGGCTCCGACGGTAAACCTACGACTGAAACGATATACTATGTGGAAACGCCAACCGTGGGAACGGCATCAACAACGTATACTCCT (135 nt) | WTGTETSTFSTETTTFTGSDGKPTTETIYYVETPTVGTASTTYTP (45 aa) |
| FLOamy | TGGACAGGAACTGAGACATCAACCTTTTCAACTGAGACTACTACTTTTACCGGTTCTGATGGTAAACCGACAACTGAAACAATAATTGTTATTAGAACTCCCACTGTCGGTACAGCTTCCACGACATATACTCCG (135 nt) | WTGTETSTFSTETTTFTGSDGKPTTETIIVIRTPTVGTASTTYTP (45 aa) |
| SHITT | GCTGGAGAAGCTGATTATACCACTACTATCACTAAGGGTAACGGTGACTTCGAAACAGACTTAGTCTCACATATTACCACGACTGACTCGGATGGCAAGCCGACTACCATCACTACTACTATTCCATTGGATGAC (135 nt) | AGEADYTTTITKGNGDFETDLVSHITTTDSDGKPTTITTTIPLDD (45 aa) |
| ALS | CGTGAGCCCCCAAATCCAACCGTAACTACTACTGAATACTGGAGTCAGTCCTACGCCACCACTACCACAGTCACGGCTCCACCAGGTGGAACAGATACTGTCATCATA (108 nt) | REPPNPTVTTTEYWSQSYATTTTVTAPPGGTDTVII (36 aa) |

Note that in the synthetic megasatellites, FLO, FLOamy and SHITT were tandemly repeated 10 times, ALS was repeated 13 times. Each motif encoded the same amino acid sequence but DNA sequences were different to facilitate synthesis. Only the first motif of the tandem repeat is shown, the complete sequence is available on request. Peptidic sequences underlined in FLO and FLOamy point to amino acid differences between the two motifs.
